# Putting pesticides on the map for pollinator research and conservation

**DOI:** 10.1101/2021.10.18.464808

**Authors:** Margaret R. Douglas, Paige Baisley, Sara Soba, Melanie Kammerer, Eric V. Lonsdorf, Christina M. Grozinger

**Author notes:** corresponding author(s): Margaret R. Douglas.

## Abstract

Wild and managed pollinators are essential to food production and the function of natural ecosystems; however, their populations are threatened by multiple stressors including pesticide use. Because pollinator species can travel hundreds to thousands of meters to forage, recent research has stressed the importance of evaluating pollinator decline at the landscape scale. However, scientists’ and conservationists’ ability to do this has been limited by a lack of accessible data on pesticide use at relevant spatial scales and in toxicological units meaningful to pollinators. Here, we synthesize information from several large, publicly available datasets on pesticide use patterns, land use, and toxicity to generate novel datasets describing pesticide use by active ingredient (kg, 1997-2017) and aggregate insecticide load (kg and honey bee lethal doses, 1997-2014) for state-crop combinations in the contiguous U.S. Furthermore, by linking pesticide datasets with land-use data in the contiguous United States, we describe a method to map pesticide indicators at spatial scales relevant to pollinator research and conservation.

## Background & Summary

With nearly 90% of flowering plant species benefiting from the services of pollinators to set seed and produce fruit, pollinators are an essential component of healthy and diverse ecosystems and contribute significantly to food production.^1–4^ However, populations of both wild and managed pollinators are facing serious challenges.^5^ Population declines have been documented in several bee and butterfly species,^6–8^ including the Eastern population of the monarch butterfly (*Danaus plexippus*), which has declined by ~80% since the mid-1990s.^9,10^ U.S. beekeepers lose around a third of their managed honey bee colonies each year.^11^ The causes of pollinator declines are multifaceted and somewhat distinct for different species, but current evidence suggests that wild bees, honey bees, and butterflies share at least two key stressors: habitat loss and pesticide exposure.^5,12,13^ Habitat loss limits the food and nesting resources available to support pollinator populations, while exposure to pesticides can kill pollinators outright or lead to sublethal effects on behavior, immunity, and reproduction.^5,12–15^ Furthermore, use of herbicides may influence pollinators indirectly by reducing the availability of their food plants.^16^

In the past decade, researchers have made significant progress in developing models to predict pollinator abundance and ecosystem services as a function of the landscape. For wild bees, the ‘Lonsdorf model’ translates land cover to abundance of nest sites and seasonal floral resources (predicted based on expert opinion), and combines this with flight ranges to derive indices of bee abundance and pollination service on each cell on a landscape;^17,18^ the model has been adapted for honey bees as well.^19^ For monarch butterflies, researchers recently developed a spatial model that simulates the annual cycle of the Eastern monarch population, identifying regions where conservation actions could enhance monarch population stability.^20^ Despite the significant value of current pollinator models based on resource availability, they could be improved by incorporating patterns of pesticide use.

There have been three main obstacles to incorporating pesticide use into landscape-scale research on pollinator health. First, although the U.S. has a substantial amount of public data on pesticide use, pesticide toxicity, and land use, these data are distributed across disparate government databases, each with idiosyncratic nomenclature and organization. Second, the mosaic of pesticide use data that are available are reported mainly at scales of counties, states, or national averages. In contrast, pollinator populations are structured at smaller spatial scales; for example bee foraging ranges are typically hundreds to thousands of meters.^21^ Finally, there are hundreds of common pesticide active ingredients that vary by many orders of magnitude in their toxicity to pollinators.^14^ Transforming pesticide use into relevant units of toxicity can help to evaluate aggregate effects.^22–25^

Here, we synthesize several large, publicly available datasets on pesticide use patterns, land use, and insecticide toxicity to generate novel datasets describing pesticide use patterns. We produced pesticide datasets by active ingredient (in kg, 1997-2017) and aggregate loading (in kg and honey bee lethal doses, 1997-2014) for state-crop combinations in the contiguous U.S. Since crop type explains much of the variation in pesticide use,^26–29^ we suggest ‘downscaling’ pesticide use by matching state-level, crop-specific pesticide use averages to land use estimated through remote sensing. We describe a method to map our pesticide use estimates to existing land use data. This methodology can readily be adapted to include more refined data that may be locally available or become available in the future, such as toxicity data for other pollinator species or site-specific information about pesticide use. Moreover, while this work is motivated primarily by the effects of pesticides on pollinators, the estimates and mapping method we describe have potential application in a wide array of settings ranging from water quality monitoring to human epidemiology.

## Methods

### Overall strategy

The aim of this project was to synthesize publicly available data on land use, pesticide use, and toxicity to generate a ‘toolkit’ of data resources enabling improved landscape-scale research on pesticide-pollinator interactions. The main outcomes are several novel datasets covering ten major crops or crop groups in each of the 48 contiguous U.S. states:

i. Average application rate (kg/ha/yr) of > 500 common pesticide active ingredients (1997-2017),
ii. Aggregate bee toxic load (honey bee lethal doses/ha/yr) of all insecticides combined (1997-2014), (Note that this dataset ends in 2014 because after that year, data on seed-applied pesticides were excluded,^30^ and these contribute significantly to bee toxic load^22^)
iii. Reclass tables relating these pesticide-use indicators to land use/land cover classes to enable the creation of maps predicting annual pesticide loading at 30-56 m resolution.

An overview of the steps, inputs, and outcomes are provided in Figure 1.

**Figure 1.**
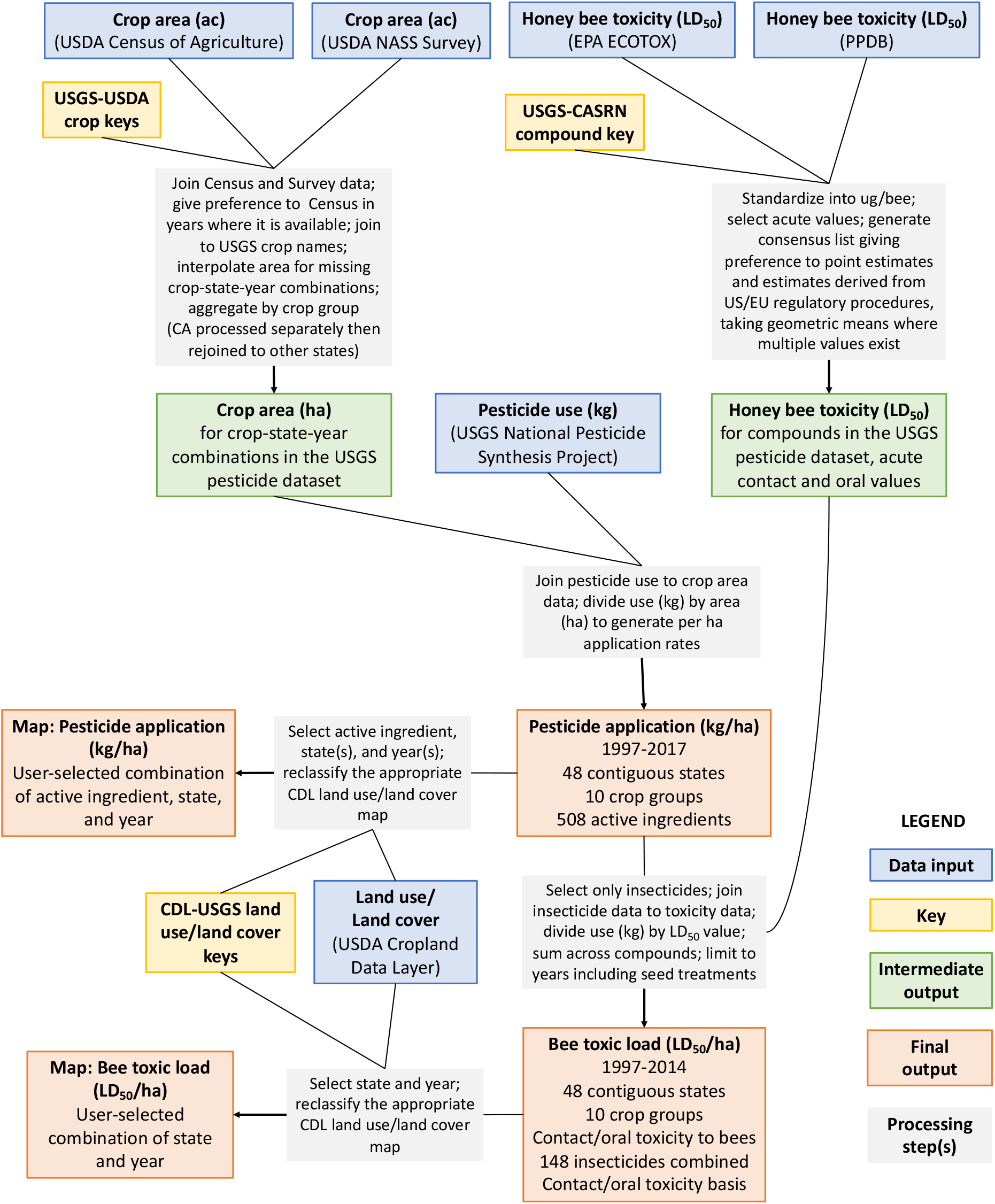
Overview of the data synthesis workflow described in this paper.

### Code availability

All data processing was performed in the R statistical language.^31^ Code is organized into a workflow that is available from the authors upon request.

### Data inputs

A summary of input datasets is provided in Table 1.

**Table 1.** Data inputs used in this study.

| Description (units) | Source(s) | Scope & resolution |
| --- | --- | --- |
| Pesticide use (kg) | USGS Pesticide National Synthesis Project<br>https://water.usgs.gov/nawqa/pnsp/usage/maps/county-level/ | > 500 pesticide active ingredients<br>Estimates for eight crop categories and the 50 contiguous states in the US<br>Annual (1992-2017) |
| Pesticide use (kg, kg/ac on treated acres; % acres treated) | USDA Agricultural Chemical Use Survey<br>https://quickstats.nass.usda.gov/ | Major crops<br>State, regional, and national resolutions<br>Annual for major crops and production areas; sporadic for minor crops (1990-2019) |
| Crop area (acres) | US Census of Agriculture<br>https://www.quickstats.nass.usda.gov/ | All crops<br>County, state, and national resolutions<br>Every five years (1997, 2002, 2007, 2012, 2017) |
|  | USDA NASS Survey<br>https://www.quickstats.nass.usda.gov/ | Major crops<br>State, regional, and national resolutions<br>Annual for major crops and production areas; sporadic for minor crops (1990-2017) |
| Honey bee acute toxicity (LD<sub>50</sub>, usually in µg/bee) | EPA ECOTOX Database<br>https://cfpub.epa.gov/ecotox/ | Wide range of toxicological measurements reported in the dataset |
|  | PPDB: Pesticides Properties Database<br>https://sitem.herts.ac.uk/aeru/ppdb/en/ | Wide range of toxicological measurements reported in the dataset |
| Land use/Land cover | USDA Cropland Data Layer<br>https://nassgeodata.gmu.edu/CropScape/ | Dozens of crop species and seven non-agricultural land cover classes for the contiguous US<br>30-56 m resolution<br>Annual (2008-2020 for entire US; some states go back to early 2000s) |

#### Pesticide data

Pesticide use data were last downloaded from the USGS National Pesticide Synthesis Project^32,33^ in June 2020. This dataset reports total kg applied of 508 common pesticide active ingredients by combinations of state, crop group, and year for the contiguous U.S. from 1992-2017. The data are derived primarily from farmer surveys conducted by a private firm (Kynetec). For California, USGS obtains data from the state’s pesticide use reporting program.^34^ USGS then aggregates and standardizes both data sources into a common national dataset that is released to the public and was used in this effort. The USGS dataset includes both a ‘high’ and a ‘low’ estimate of pesticide use, varying based on the treatment of missing values in the source data.^33^ We used the ‘low’ estimate throughout, but assess the influence of this choice on the resulting estimates (see Technical Validation).

#### Crop area data

To translate pesticide use estimates into average application rates, it was necessary to divide total kg of pesticide applied by the land area to which it was potentially applied. Crop area data were last downloaded from the Quick Stats Database of the USDA^35^ in May 2020, using data files downloaded from the ‘developer’ page. This USDA dataset contains crop acreage estimates generated from two sources: the Census of Agriculture (Census), which is comprehensive but conducted only once every five years^36^ and the crop survey conducted by the National Agricultural Statistics Service (NASS), which is an annual survey based on a representative sample of farmers in major production regions for a more limited subset of crops.^37^

#### Honey bee toxicity data

Translating insecticide application rates into estimates of bee toxic load (honey bee lethal doses/ha/yr) required toxicity values for each insecticide active ingredient in the USGS dataset. We used LD_50_ values for the honey bee (*Apis mellifera*) because this is the standard terrestrial insect species used in regulatory procedures, and so has the most comprehensive data available. This species is also of particular concern as an important provider of pollination services to agriculture. As previously reported,^22^ the LD_50_ values were derived from two sources, the ECOTOX database^38^ of the U.S. Environmental Protection Agency (US-EPA), and the Pesticide Properties Database (PPDB).^39^ ECOTOX was queried in July 2017, by searching for all LD_50_ values for the honey bee (*Apis mellifera*) that were generated under laboratory conditions. Acute contact and oral LD_50_ values for the honey bee were recorded manually from the PPDB in June 2018.

#### Land cover data

Mapping pesticides to the landscape requires land use/land cover data indicating where crops are grown. We used the USDA Cropland Data Layer (CDL),^40^ a land cover dataset at 30-56 m resolution produced through remote sensing. This dataset is available from 2008-2020 for states in the contiguous U.S., with some states (primarily in the Midwest and Mid-South) available back to the early 2000s.

### Data preparation

#### Relating datasets

A major challenge in this data synthesis effort was relating the various data sources to each other, given that each dataset has unique nomenclature and organization. We created the following keys (summarized in Table 2) to facilitate joining datasets:

i. *USGS-USDA crop keys* – Using documentation and metadata associated with the USGS pesticide dataset,^33,41,42^ we created keys relating the USGS surveyed crop names (‘ePest’ crops) and the ten USGS crop categories to the large number of corresponding crop acreage data items in the Census and NASS datasets. For annual crops and hay crops we used ‘harvested acres,’ and for tree crops we used ‘acres bearing & non-bearing.’ These choices were made to maximize data availability and to correspond as closely as possible to the crop acreage from which the pesticide data were derived.^33^ A separate key was developed for California because California pesticide data derives from different source data and covers a larger range of crops.
ii. *USGS-CASRN compound key* – Using USGS documentation as well as background information on pesticide active ingredients,^39,43^ we generated keys relating USGS active ingredient names to chemical abstracts service (CAS) registry numbers to facilitate matching compounds to the ECOTOX and PPDB databases.
iii. *USGS compound-category key* – In this key we classified active ingredients into major groups (insecticides, fungicides, nematicides, etc.) and into mode-of-action classes on the basis of information from pesticide databases and resistance action committees.^43–46^
iv. *USGS-USDA compound key* – To facilitate our data validation effort, we generated a key relating USGS compound names to USDA compound names, on the basis of information from several pesticide databases.^39,43^
v. *USGS-CDL land use-land cover keys* – Using documentation from the USGS pesticide dataset describing the crop composition of each of the ten crop categories,^33^ we created a key that matches these categories to land cover classes in the CDL. A separate key was developed for California given the differences in surveyed crops in this state, noted above.

**Table 2.** Keys generated to relate datasets

| Key name | Purpose | File name(s) |
| --- | --- | --- |
| USGS-USDA<br>crop keys | Relate USDA crop names/data items to surveyed crops and crop groups in the USGS pesticide dataset | crop_key_summary.csv<br>crop_key_summary_CA.csv |
| USGS-CASRN<br>compound key | Relate compound names in the USGS pesticide dataset to CAS Registry Numbers for comparison to other datasets | USGS_Pesticide-CASRN.csv |
| USGS compound -<br>category key | Categorize compounds by type (insecticide, fungicide, herbicide) and mode-of-action group | USGS_Pesticide_Category.csv |
| USGS-USDA<br>compound key | Relate compound names in the USGS pesticide dataset to pesticide-related data items in USDA Agricultural Chemical Use Survey. | usda_usgs_pesticide_names.csv |
| USGS-CDL land<br>use-land cover keys | Relate crop groups from the USGS pesticide dataset to land cover classes in the USDA Cropland Data Layer | cdl_reclass.csv<br>cdl_reclass_CA.csv |

#### Processing crop area data

Because of differences in the crops included in pesticide use estimates, crop acreage data were processed separately for California and for all other states, and then re-joined, as follows: Acreage data were first filtered to include only data at the state level, reporting total annual acreage for states in the contiguous U.S. after 1996. Acreage data were joined to the appropriate USGS-USDA crop key and only those crops represented in the pesticide dataset were retained. We then generated an acreage dataset with single rows for each combination of crop, state, and year using data from the Census when available (1997, 2002, 2007, 2012, 2017), data from NASS in non-census years, and linear interpolation to fill in remaining missing values. This process was repeated for California, using acreage data for only that state in combination with the CA crop key. Finally, acreage data in the two datasets were recombined, converted to hectares, and summed by USGS crop group.

#### Processing honey bee toxicity data

Processing for the honey bee toxicity data has been described in detail elsewhere.^22^ Briefly, toxicity values from ECOTOX were categorized as contact, oral, or other and standardized where possible into μg/bee. Records were retained if they represented acute exposure (4 days or less) for adult bees representing contact or oral LD_50_ values in μg/bee. To generate a consensus list of contact and oral LD_50_ values for all insecticides reported in the USGS dataset, we gave preference to point estimates and estimates generated through U.S. or E.U. regulatory procedures, taking a geometric mean if multiple such estimates were available. Unbounded estimates (“greater than” or “less than” some value) were only used when point estimates were unavailable, using the minimum (for “less than”) or the maximum (for “greater than”). If values for a compound were unavailable in both datasets, we used the median toxicity value for the insecticide mode-of-action group. And finally, in rare cases (n = 1/148 compounds for contact toxicity and 8/148 compounds for oral toxicity) we were still left without a toxicity estimate for a particular insecticide. In those cases, we used the median value for all insecticides.

### Data synthesis

#### Compound-specific application rates for state-crop-year combinations

USGS data on pesticide application were joined to data on crop area. Average pesticide application rates were calculated by dividing kg applied by crop area (ha) for each combination of compound, state, and year.

#### Aggregate insecticide application rates for state-crop-year combinations

The dataset from the previous step was filtered to include only insecticides, and then joined to LD_50_ data by compound name. Bee toxic load associated with each insecticide active ingredient was calculated by dividing the application rate by the contact or oral LD_50_ value (μg/bee) to generate a number of lethal doses applied per unit area. These values were then summed across compounds to generate estimates of kg and bee toxic load per ha for combinations of crop, state, and year. Missing values were estimated using linear interpolation from other years for the same state-crop combination, where possible. We focused the bee toxic load estimates on insecticides because *i*) insecticides are used to kill insects, and so tend to have higher acute toxicity toward insects than other types of pesticides, and *ii*) honey bee toxicity data for fungicides and herbicides are inconsistently available. This dataset ends in 2014 because after that year seed-applied pesticides were excluded from the source data,^30^ and they constitute a major contribution to bee toxic load.^22^

#### Reclassification tables

To generate reclassification tables for the CDL, the pesticide datasets described above were joined by crop to CDL land use categories. The output of these processes was a set of reclassification tables for bee toxic load and each individual compound for all combinations of state, year, and land use category.

Of the 131 land use categories in the CDL, 16 represent two crops grown sequentially in the same year (double crops, found on ~2% of U.S. cropland in 2012^47^), which required a modified accounting in our workflow. Pesticide use practices on double crops are not well described, but one study suggested that pesticide expenditures on soybean grown after wheat were similar to pesticide expenditures in soybean grown alone.^48^ Therefore, we assumed that pesticide use on double crops would be additive (e.g. for a wheat-soybean double crop, the annual pesticide use estimate was generated by summing pesticide use associated with wheat and soybean).

Missing values in the reclassification tables resulted from several distinct issues. Some values were missing because a particular crop was not included in the underlying pesticide use survey (e.g. oats was not included in the Kynetec survey), or because the land use category was not a crop at all (e.g. deciduous forest). These two issues were indicated with values of ‘1’ in columns called ‘unsurveyed’ and ‘noncrop,’ respectively. For double crops, a value of 0.5 in the ‘unsurveyed’ column indicates that one of the crops was surveyed and the other was not. For compound-specific datasets, missing values may reflect that a given compound is not used in a state-crop-year combination. For the aggregate insecticide dataset, even after interpolation there were some missing values, usually when a state had very little area of a particular crop.

Finally, missing data for double crops were treated slightly differently in the aggregate vs. compound-specific reclassification tables. For the aggregate insecticide dataset, estimates for double crops were only included if estimates were available for both crops; otherwise the value was reported as missing. For the compound-specific datasets, estimates for double crops were included if there was an estimate for at least one of the crops, since specific compounds may be used in one crop but not another.

### Data Records

Data files generated in this work are available from the authors upon request. A summary of output data files is provided in Table 3.

**Table 3.** Data outputs generated by this study

| Output name<br>(units) | Scope & resolution | File name(s) |
| --- | --- | --- |
| Honey bee toxicity for compounds in the USGS pesticide dataset (µg/bee) | 148 insecticide active ingredients<br>Contact + oral acute toxicity | ld50_usgs_complete_20200609.csv |
| Crop area for crops represented in the USGS pesticide dataset (ha) | 1997-2017 (annual)<br>48 contiguous states<br>87 crops (140 in California)<br>10 crop groups | hectares_state_usda_usgs_20200404.csv<br>hectares_CA_usda_usgs_20200404.csv |
| Pesticide application (kg/ha/yr) | 1997-2017 (annual)<br>48 contiguous states<br>10 crop groups<br>508 active ingredients | bee_tox_index_state_yr_cmpd_20200609.csv |
| Insecticides aggregated (kg/ha/yr + honey bee lethal doses/ha/yr) | 1997-2014 (annual)<br>48 contiguous states<br>10 crop groups<br>148 insecticides combined<br>Contact/oral toxicity basis | bee_tox_index_state_yr_cat_20210625.csv |
| Cropland Data Layer reclass master files - pesticide application (kg/ha/yr) | 1997-2017 (annual)<br>48 contiguous states<br>131 land use/land cover classes<br>508 active ingredients | beetox_cmpd_cdl_reclass_20210607<br>[separate file for each active ingredient of the form COMPOUND.csv] |
| Cropland Data Layer reclass master file – insecticides aggregated (kg/ha/yr + honey bee lethal doses/ha/yr) | 1997-2014 (annual)<br>48 contiguous states<br>131 land use/land cover classes<br>148 insecticides combined<br>Contact/oral toxicity basis | beetox_I_cdl_reclass.20210625.csv |

### Technical Validation

#### Validity of pesticide use estimates

To assess the validity of the dataset, we compared our estimates for specific compounds to those available from the USDA’s Agricultural Chemical Use Program^49^ for the period 1997-2017. While this dataset is not as comprehensive as the USGS dataset in terms of its coverage of states, years, crops, and active ingredients, the values that are available can be compared to USGS values for total kg applied and estimates of kg/ha/yr. An important limitation of this comparison is that the USDA dataset does not include seed-applied pesticides, affecting the estimation of certain fungicide and insecticide active ingredients.^30^

For combinations of state, crop, and active ingredient, the USDA survey reports several values that are relevant to this effort: total amount (kg) of pesticide applied, percent area treated, and application rate (kg/ha) on treated hectares. Application rates across the two datasets are not directly comparable because USDA reports the rate (kg/ac) applied on treated acres, whereas our dataset estimates the rate (kg/ha) averaged across all crop area in a state. To test the agreement between our estimates and the USDA estimates we compared:

i. Total amount applied (kg),
ii. Application rate (kg/ha) calculated by dividing the USDA total kg by our estimate of crop area (ha) (Method 1),
iii. Application rate (kg/ha) derived indirectly by multiplying the USDA average application rate on treated acres by the percent of area treated (Method 2).

We focused on major field crops (corn, soy, wheat, cotton, rice) because they are the most frequently surveyed by USDA and have the closest one-to-one match between datasets. We suspect that estimates for fungicide and insecticide use may be more precise in high-value fruit and vegetable crops, where these products are more widely and heavily used.^26,28^ However, fruit and vegetable crops could not be included in the validation because estimates are incommensurable across the two datasets: USGS reports pesticide use for crop groups like ‘orchards and grapes’ while USDA reports pesticide use on a small subset of individual crops in major production states (e.g. apples in Washington).

For each set of comparisons, we calculated and visualized relative percent difference between the two estimates, visualized the correlation between estimates, and calculated Spearman’s and Pearson’s correlation coefficients for each major pesticide class (fungicides, herbicides, insecticides).

Estimates for total amount applied and per-ha application rate were strongly correlated for fungicides and herbicides, with Pearson’s correlation coefficients > 0.85 for all comparisons (Table 4, Figure 2). Estimates for insecticides were well correlated on the basis of ranks (Spearman’s *rho* = 0.85) but only weakly linearly related (Pearson’s *r* = 0.20-0.38, Table 4, Figure 2). This pattern was driven by malathion in cotton, which had very low estimates in the USGS dataset and fairly high estimates in the USDA dataset, for reasons unknown. Once these outliers were removed (*n* = 30 out of 1600+ observations), estimates for insecticides were well correlated for all three comparisons (Pearson’s *r* > 0.75).

**Table 4.** Statistical results from cross-validation comparing USDA pesticide use estimates to the estimates generated in this study from USGS pesticide use data and USDA crop area data. For application rate, Method 1 (M1) compared our estimate to one calculated by dividing the USDA total kg by our estimate of crop area (ha), and Method 2 (M2) compared our estimate to one calculated by multiplying the USDA average application rate on treated hectares by the percent of area treated. Relative difference = [(USGS – USDA)/average of the two values] × 100.

| Category | Comparison | N<br>(# state-crop-year-AI) | Relative difference (%): |  |  |  | Spearman's<br>*rho* | Pearson's<br>*r* |
| --- | --- | --- | --- | --- | --- | --- | --- | --- |
|  |  |  | Median | 25<sup>th</sup><br>percentile | 75<sup>th</sup><br>percentile | Wilcoxon<br>test<br>*p* |  |  |
| Fungicides | kg applied | 501 | 4 | -51 | 54 | 0.59 | 0.79 | 0.86 |
|  | kg/ha (M1) | 501 | 4 | -51 | 54 | 0.59 | 0.71 | 0.95 |
|  | kg/ha (M2) | 502 | 2 | -56 | 56 | 0.99 | 0.71 | 0.94 |
| Herbicides | kg applied | 6762 | 2 | -38 | 40 | 0.45 | 0.94 | 0.97 |
|  | kg/ha (M1) | 6762 | 2 | -38 | 40 | 0.45 | 0.94 | 0.95 |
|  | kg/ha (M2) | 6891 | 3 | -39 | 42 | 0.04 | 0.94 | 0.94 |
| Insecticides | kg applied | 1699 | -14 | -69 | 43 | < 0.01 | 0.85 | 0.20 |
|  | kg/ha (M1) | 1699 | -14 | -69 | 43 | < 0.01 | 0.85 | 0.36 |
|  | kg/ha (M2) | 1712 | -10 | -65 | 47 | < 0.01 | 0.85 | 0.38 |
| Insecticides* | kg applied | 1669 | -12 | -66 | 45 | < 0.01 | 0.87 | 0.84 |
|  | kg/ha (M1) | 1669 | -12 | -66 | 45 | < 0.01 | 0.86 | 0.77 |
|  | kg/ha (M2) | 1682 | -8 | -62 | 48 | < 0.01 | 0.87 | 0.78 |
| All active ingredients | kg applied | 8962 | -0.1 | -45 | 41 | 0.01 | 0.92 | 0.93 |
|  | kg/ha (M1) | 8962 | -0.1 | -45 | 41 | 0.01 | 0.92 | 0.83 |
|  | kg/ha (M2) | 9105 | 1 | -45 | 44 | 0.30 | 0.92 | 0.83 |

\*Recalculated after removing malathion use in cotton

**Figure 2.**
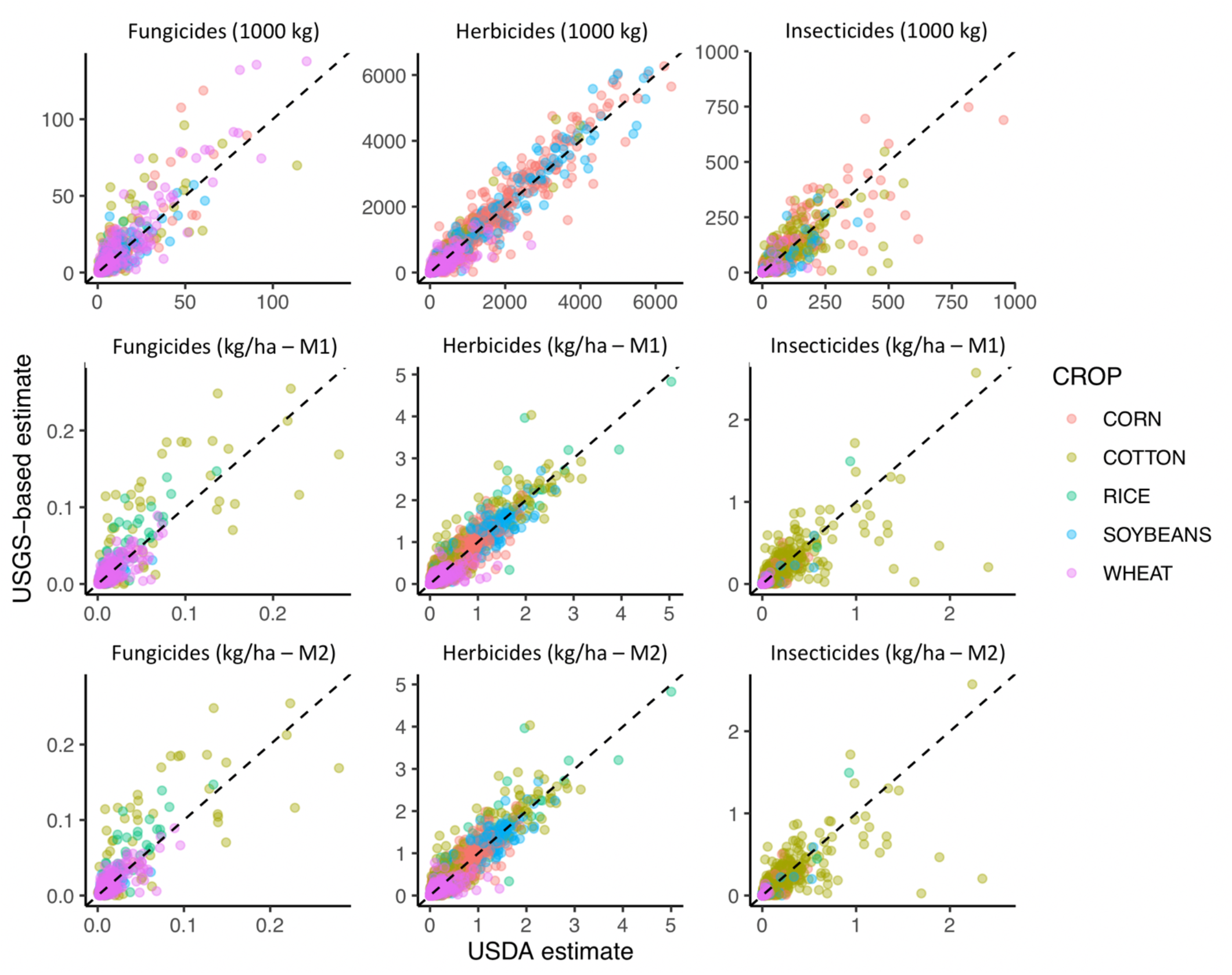
Correlation between USDA pesticide use estimates and novel estimates generated from USGS data reported in this paper. Dotted black line shows USDA = USGS. For application rate, Method 1 (M1) compared our estimate to one calculated by dividing the USDA total kg by our estimate of crop area (ha), and Method 2 (M2) compared our estimate to one calculated by multiplying the USDA average application rate on treated hectares by the percent of area treated. Each point represents a combination of state, crop, year, and active ingredient. Outliers (malathion in cotton [n = 30] and copper sulfate in rice [n = 2]) were removed before plotting.

Among pesticide categories, the median relative difference between USGS-based datasets and USDA estimates was smallest for herbicides and largest for insecticides (Table 4). For fungicides, the median relative difference did not differ significantly from zero for any of the three comparisons, indicating a lack of consistent over- or under-estimation. For herbicides, median relative difference was not significantly different from zero for total kg or application rate (Method 1). However, it was significantly positive for application rate (Method 2), indicating a tendency for the USGS-based estimate to be slightly larger than the USDA-based estimate (median difference of 3%). For insecticides, median relative difference was significantly less than zero for all comparisons (−10-14%), indicating that the USGS-based estimates are conservative compared to the USDA estimates. Median relative difference decreased slightly when malathion outliers were removed (−8-12%), but were still significantly less than zero. This result may derive from using the USGS ‘low’ pesticide use values (see Sensitivity analyses).

Importantly, relative difference between estimates declined with percent of crop area treated (Figure 3), suggesting that estimates are more precise for active ingredients that are widely used. This pattern may help to explain why median relative difference between datasets was smallest for herbicides and largest for insecticides, given that field crops formed the basis of the validation. In field crops, herbicide use is very widespread while use of traditional insecticides and fungicides is more limited.^29^ Since the mid-2000’s the most widely applied insecticides in field crops have been neonicotinoid seed treatments,^30,50^ which are excluded from the USDA survey and by extension, this comparison. Previous work^50^ suggests that USGS estimates of neonicotinoid use in major field crops are consistent with independent estimates.

**Figure 3.**
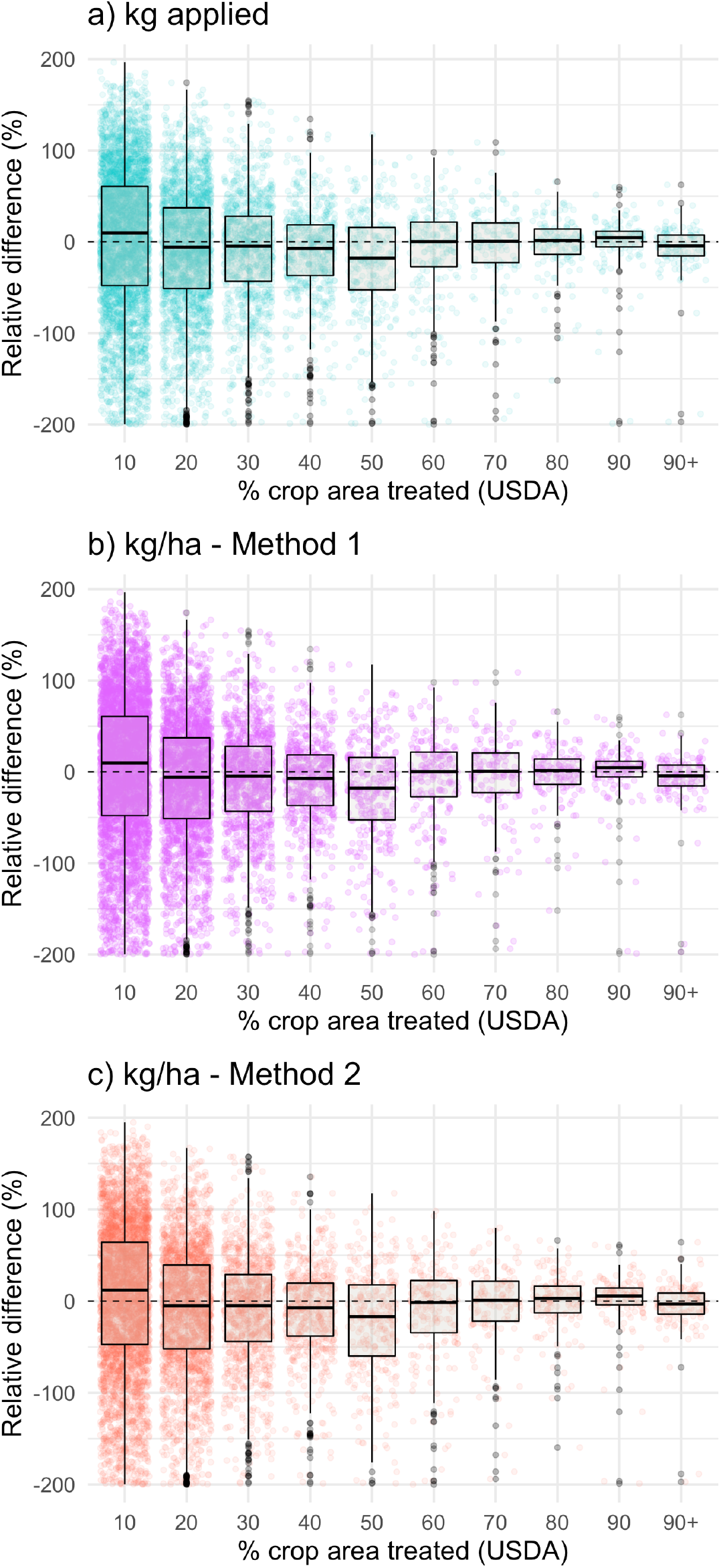
Relationships between the percent of cropland treated with a pesticide and the relative difference between the USDA pesticide use estimates and novel estimates generated from USGS data reported in this paper. Each point represents a combination of state, crop, year, and active ingredient. Relative difference = [(USGS – USDA)/average of the two values] × 100.

#### Geographic coverage

To characterize the geographic coverage of our datasets, we calculated the percent of total land area and agricultural area in each state that is included in the underlying pesticide use survey. To do this, we downloaded the CDL for all states in even years, calculated the area in each CDL category, joined the data to the CDL-USGS land use/land cover key, and aggregated the area depending on whether it was surveyed, unsurveyed, or non-agricultural. In most states, the survey covers > 95% of agricultural land (> 80% of agricultural land in all states, Figure 4). States on the lower end of this range have significant area of regionally important yet unsurveyed crops (e.g. blueberries in Maine, cranberries in Massachusetts, grass seed in Oregon). Double crops contributed < 5% to agricultural area in most states, except North Carolina, Maryland, and Delaware, where they comprised 5-23% of agricultural area. As expected, survey coverage of total land area was variable among states, ranging from < 10% in states dominated by forest (e.g. New Hampshire) or shrubland (e.g. Nevada), to > 80% in states with abundant cropland (e.g. Iowa).

**Figure 4.**
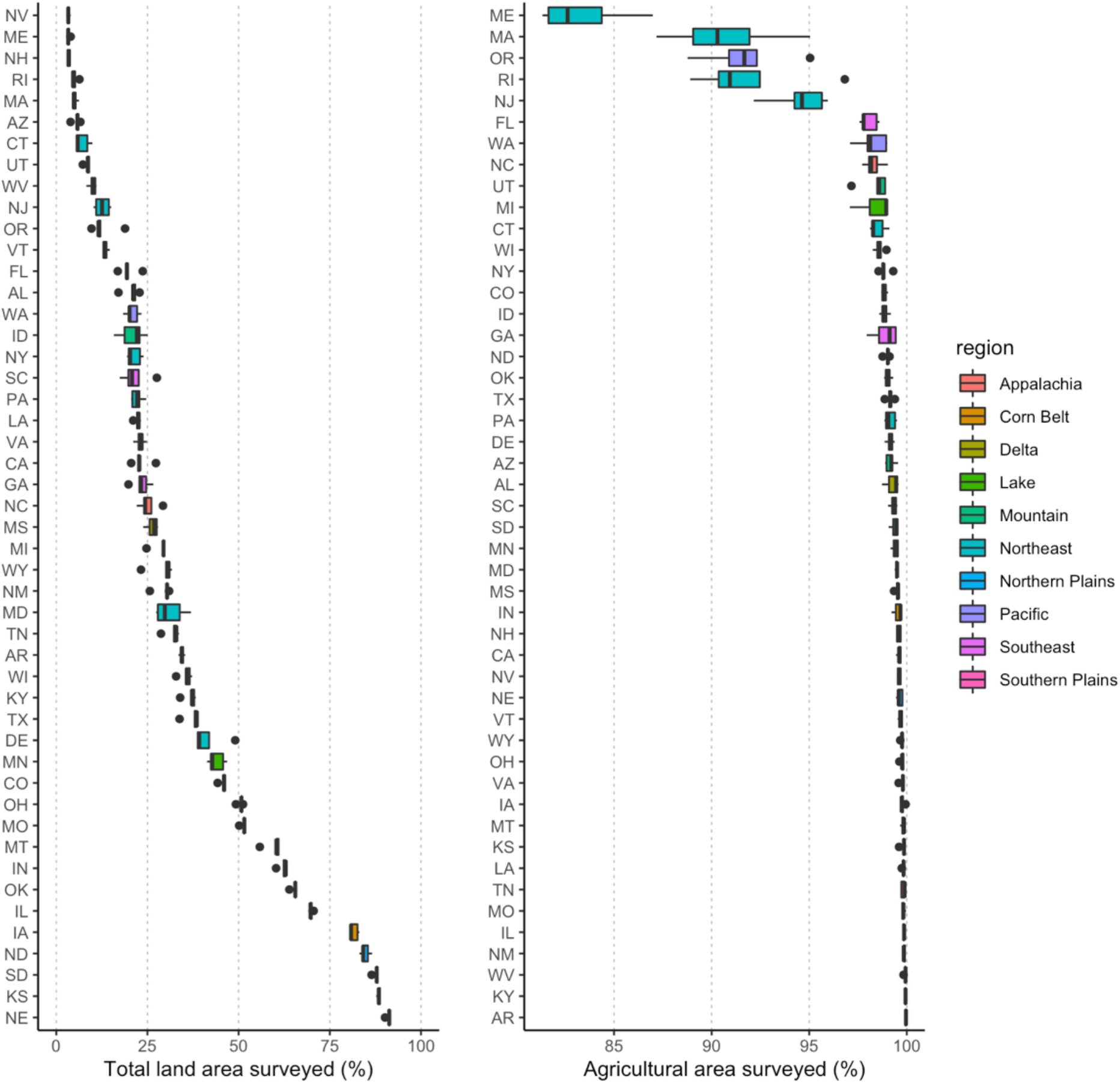
Pesticide survey coverage for the 48 contiguous states in even years from 2008-2016. (left) Percent of total land area represented by surveyed land uses and (right) percent of cropland represented by surveyed land uses (where ‘surveyed’ means included in the underlying pesticide use survey).

#### Sensitivity analyses

We conducted several additional analyses to quantify the sensitivity of our results to the methods employed in generating the dataset. To test the influence of using the low vs. the high USGS pesticide estimate, we regenerated the dataset using the high estimate and compared validation metrics. Consistent with previous analyses,^42^ the high estimate from USGS tended to exceed the USDA value, with median relative differences ~30% for herbicides, 10-12% for fungicides, and 6-10% for insecticides. The low estimate was closer to the USDA value except in the case of insecticides, in which the low estimate underestimated the USDA value to a slightly greater degree than the high estimate overshot. However, correlations between the two estimates were similarly strong regardless of whether the high or low estimate was used (Pearson’s *r* > 0.75 for all comparisons), suggesting that relative patterns in the data are robust.

To determine the potential uncertainty introduced by interpolating crop area when it was missing, we investigated the amount of total area contributed by interpolated values across the dataset. Across all years combined, interpolated area was a small percentage of most crops or crop groups (< 4% of total area), with the exceptions of ‘orchards and grapes’ (76%), ‘pasture and hay’ (76%), and ‘vegetables and fruit’ (36%). Estimates for land area in these groups relied heavily on interpolation between Census years, because these land uses are infrequently surveyed apart from the Census. We are confident in the interpolated values because they are mostly for perennial crops, so it is unlikely that acreage would change substantially from year to year. That said, for the most accurate estimates, we recommend using the pesticide datasets matching Census years (1997, 2002, 2007, 2012, 2017).

We also investigated the influence of interpolated values for aggregate insecticide indicators. While interpolated values occupied a significant proportion of surveyed agricultural area (59%), they contributed < 1.5% to total insecticide load in kg or honey bee LD_50_s. This pattern was driven by extensive interpolation for ‘pasture and hay,’ a land use category that occupies significant land area but typically has very low insecticide inputs. Conversely, land use categories with heavy insecticide inputs (e.g. cotton, ‘vegetables and fruits’, ‘orchards and grapes’) had little to no interpolation.

### Usage Notes

#### R functions to generate maps

One of the main goals of this project was to develop a methodology to generate maps of predicted pesticide use and bee toxic load. We provide two R functions (available in the ‘beecoSp’ package, https://github.com/land-4-bees/beecoSp) to reclassify land cover into predicted pesticide application rate (kg/ha) for combinations of active ingredients, states, and years, or to reclassify land cover into predicted bee toxic load (honey bee LD_50_s/ha) for all insecticides combined. The ‘reclasstables’ function facilitates the creation of reclass tables for particular state-year combinations from the master files provided here, while the ‘CDL_reclass’ function uses such a reclass table to reclassify a land use raster, with the option to calculate the mean of the new raster. The inputs include an appropriate land cover raster file (from the CDL) and an appropriate master reclassification file from those accompanying this paper. Examples of the input file and output maps for a particular active ingredient and for bee toxic load are shown in Figure 5. Alternatively, for users interested in performing reclassification in another GIS program, reclassification tables may be generated from the datasets produced here or downloaded from the website described below.

**Figure 5.**
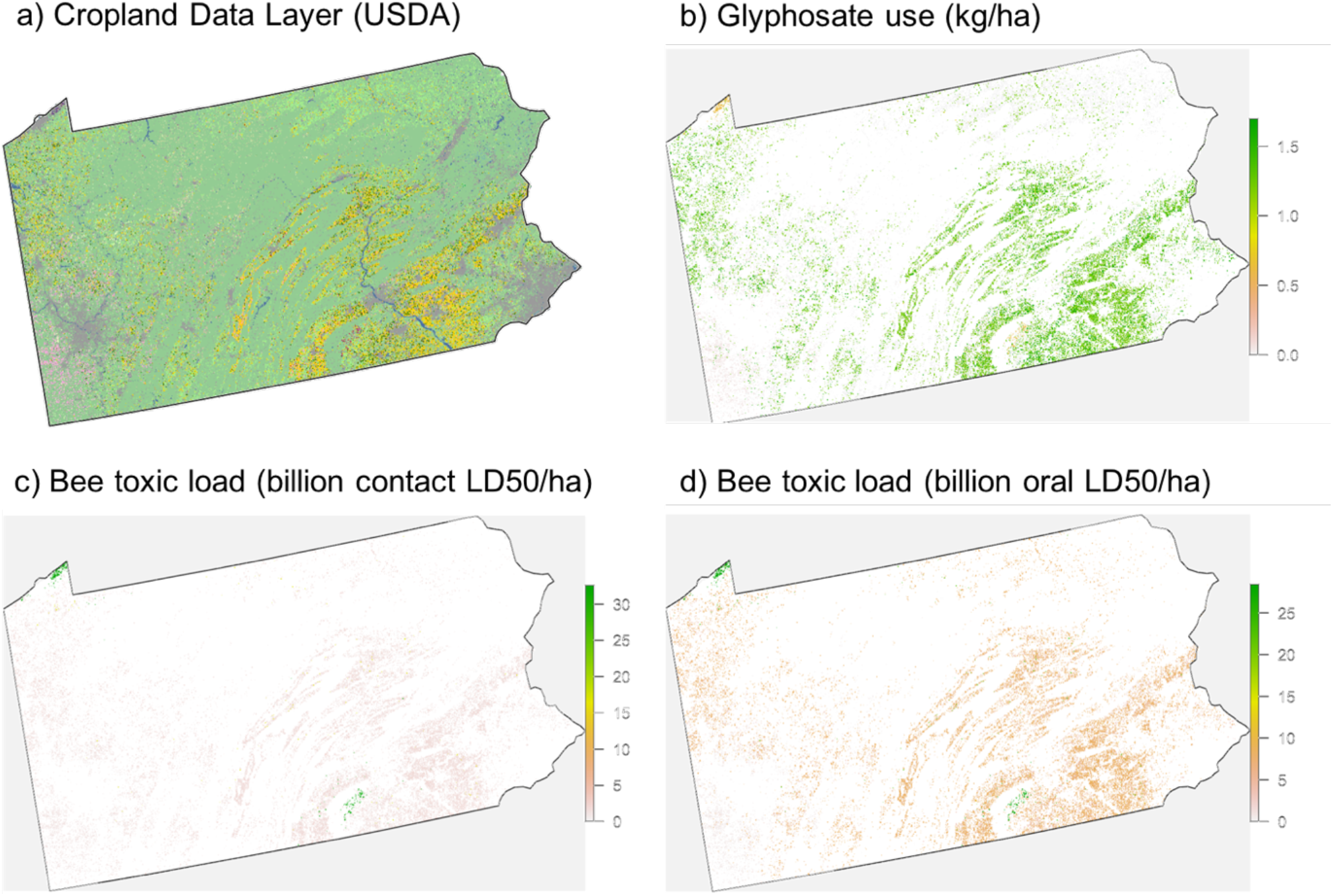
Maps illustrating the conversion of land cover to predicted landscape loading of agricultural pesticide use. Maps are shown for Pennsylvania in 2012 representing (a) input raster from the Cropland Data Layer, (b) output raster illustrating a single compound (the herbicide glyphosate), (c,d) output rasters illustrating bee toxic load for all insecticides combined, on a contact- and oral-toxicity basis (respectively).

#### Insecticide Explorer web application

To enable users to explore insecticide use patterns and easily download reclassification tables for bee toxic load, we created an interactive website. The website allows users to generate graphs describing trends in national and state-level total and per ha insecticide use (in kg and honey bee lethal doses per year) from 1997-2014 (‘Explore’ tab). There is also a section of the website allowing download of reclassification tables for bee toxic load for particular state-year combinations (‘Data’ tab). The application can be accessed at the following URL: https://insecticideexplorer.shinyapps.io/insecticideexplorer/.

#### Limitations and possible adjustments

Users should be aware of several limitations of these data. First, the pesticide use estimates include only pesticides applied to agricultural land, and so do not account for pesticide applications on other land uses, i.e. urban and unmanaged areas. Similarly, for minor crops (e.g. fruits and vegetables) the estimates of pesticide use represent averages for crop groups rather than crop-specific estimates, and some crops were not surveyed and so are not included. If users have access to independent pesticide use estimates for non-agricultural land covers, unsurveyed crops, or crops included in larger categories, the reclassification tables could be adjusted or supplemented to better represent pesticide use on those land cover classes.

Second, these estimates represent crop-state-year pesticide use averages, not field-specific application. Although crop groups, states, and years capture significant variation in pesticide use,^26–29^ there will be significant local variation not reflected in these estimates. Likewise, the resulting maps are constrained by the quality of the underlying land cover data, which are most accurate for large-acreage crops and major production regions and least accurate for minor crops and minor production regions.^51,52^ These estimates are therefore most suited to applications in which the goal is to characterize or compare pesticide use patterns over broad areas (e.g. bee foraging ranges, watersheds), not applications in which field-scale accuracy is paramount. And finally, the temporal dimension of these estimates is limited by the source data, which are reported annually, and complete only through 2014, after which seed-applied pesticides are excluded. The annual nature of the data constrains their use in mechanistic risk assessments, which require more specific information about site, method, and time of pesticide application. The exclusion of seed treatments after 2014 is problematic because seed-applied pesticides are increasingly prevalent in U.S. agriculture^30^ and seed-applied neonicotinoids represent the most important contributor to the bee toxic load calculated based on oral toxicity.^22,23^ We encourage users to investigate temporal patterns in use estimates for their target area to determine the most appropriate course to take for years after 2014. In some cases, projecting earlier use rates to later years – while clearly imperfect – may be the best available option until more valid public data becomes available.

There are also important limitations related to the ‘bee toxic load’ estimate for all insecticides combined. Most obviously, because they are based on honey bee LD_50_ units, these estimates are most relevant to honey bees and closely related species. The literature suggests that broad patterns of toxicity are similar between honey bees and other bee species,^53^ but there is significant variation among species, which is more pronounced when more distant insect taxa are considered.^54,55^ If a quantitative estimate of lethal doses for some other bee species is desired, at minimum, users should consider adjusting for body weight (honey bee workers weigh ~120 mg each^56^). Also, because these estimates are based on acute toxicity to adult honey bees, they do not reflect chronic toxicity or effects on other life stages such as larvae. It is up to the user to decide if toxic load on a contact- or oral-toxicity basis, or application rate of specific active ingredients, is most appropriate for a given application. Finally, because we summed across active ingredients, our estimate does not account for potential synergy between them. And lastly, these estimates represent loading of pesticides to the landscape but not persistence or movement in the environment. These data could be combined with additional information to model fate in the environment and exposure of recipient organisms.

## Acknowledgements

This work was supported by the National Socio-Environmental Synthesis Center (SESYNC) under funding received from the National Science Foundation (DBI-1639145) and by grants from USDA-NIFA-AFRI (#2018-67013-27538), the Foundation for Food and Agricultural Research (#549032), and a cooperative agreement from USDA-ERS (#58-30000-5-0037). We thank Nancy Baker and Wes Stone for help interpreting USGS metadata for the pesticide source data, and members of the ‘Pesticides and Pollinators’ working group, who contributed valuable suggestions during development of this project. We thank Karan Shakya and David Smith for assistance creating several keys.

## Author contributions

Margaret R. Douglas devised the data synthesis strategy, drafted most of the data processing pipeline, and drafted the manuscript.

Paige Baisley assisted with the technical validation.

Sara Soba created the ‘Insecticide Explorer’ web application.

Melanie Kammerer assisted with code and function development.

Eric V. Lonsdorf contributed to the pipeline logic and strategy for downscaling the estimates.

Christina M. Grozinger worked with Douglas on the conceptual development of the overall project goals and pipeline.

## Competing interests

The authors have no competing interests to declare.

## Notes

### Competing Interest Statement

The authors have declared no competing interest.

